# Detection of orthologous genes with expression shifts linked to nickel hyperaccumulation across Eudicots

**DOI:** 10.1101/2022.09.28.509953

**Authors:** Mélina Gallopin, Christine Drevet, Vanesa S. Garcia de la Torre, Sarah Jelassi, Marie Michel, Claire Ducos, Cédric Saule, Clarisse Majorel, Valérie Burtet-Sarramegna, Yohan Pillon, Paul Bastide, Olivier Lespinet, Sylvain Merlot

## Abstract

The remarkable capacity of plants to tolerate and accumulate tremendous amount of nickel is a complex adaptative trait that appeared independently in more than 700 species distributed in about fifty families. Nickel hyperaccumulation is thus proposed as a model to investigate the evolution of complex traits in plants. However, the mechanisms involved in nickel hyperaccumulation are still poorly understood in part because comparative transcriptomic analyses struggle to identify genes linked to this trait from a wide diversity of species. In this work, we have implemented a methodology based on the quantification of the expression of orthologous groups and phylogenetic comparative methods to identify genes which expression is correlated to the nickel hyperaccumulation trait. More precisely, we performed *de novo* transcriptome assembly and reads quantification for each species on its own transcriptome using available RNA-Seq datasets from 15 nickel hyperaccumulator and non-accumulator species. Assembled contigs were associated to orthologous groups built using proteomes predicted from completed plant genome sequences. We then analyzed the transcription profiles of 5953 orthologous groups from distant species using a phylogenetic ANOVA. We identified 31 orthologous groups with an expression shift associated with nickel hyperaccumulation. These orthologous groups correspond to genes that have been previously implicated in nickel accumulation, and to new candidates involved in this trait. We thus believe that this method can be successfully applied to identify genes linked to other complex traits from a wide diversity of species.

## Introduction

Comparative biology is a fundamental strategy to study the evolution of complex developmental and physiological traits. The development of the RNA-Seq technology has opened the possibility to perform comparative studies at the molecular level in a wide diversity of plant species (Leebens-Mack *et al*., 2019). In plants, RNA-Seq data have been used to compare distant species for phylogenetic tree inference and to study the evolution of complex traits such as the type of photosynthesis or the capacity to establish symbioses (Jiao *et al*., 2011; Wickett *et al*., 2014; Yang *et al*., 2015; Heyduk *et al*., 2019; Radhakrishnan *et al*., 2020; Rich *et al*., 2021). In most of these studies, RNA-Seq data are used to reveal gene loss, gene duplication and genetic variations linked to a specific trait in different phylae. However, because of the difficulty to use a unique sequence as reference, comparative studies rarely take full advantage of the quantitative information enclosed in RNA-Seq data to compare gene expression between distant species and identify genes which expression is linked to a particular trait (Roux *et al*., 2015; Voelckel *et al*., 2017; García de la Torre *et al*., 2021; Rich *et al*., 2021).

Metal hyperaccumulation represents an interesting case study to identify genes linked to a complex adaptative trait over a wide diversity of plant species (Manara *et al*., 2020). Metal hyperaccumulation is defined as the capacity of plant species to accumulate in their leaves a high concentration of metal, such as nickel, manganese, zinc or cadmium, that is normally toxic for the vast majority of plants (van der Ent *et al*., 2013). Today, about 700 plant species are known to hyperaccumulate metals but the large majority (*ie* 500 species) hyperaccumulates nickel (Reeves *et al*., 2018). Nickel hyperaccumulators are distributed in about 50 families among more than 300 families of dicotyledon plants suggesting that the nickel hyperaccumulation trait appeared independently in several clades along plant evolution (Krämer, 2010; Cappa and Pilon-Smits, 2014). Comparative transcriptomic analysis of zinc and cadmium hyperaccumulators and non-accumulator species of the Brassicaceae family has first revealed that metal hyperaccumulation is linked to the high and constitutive expression of several genes involved in metal transport and homeostasis (Hammond *et al*., 2006; Weber *et al*., 2006; Hanikenne *et al*., 2008; Halimaa *et al*., 2014). Our knowledge of the molecular mechanisms involved in nickel hyperaccumulation is still limited but, as for the hyperaccumulation of zinc and cadmium, the hyperaccumulation of nickel likely evolved from the high and constitutive expression of genes involved in metal homeostasis. Halimaa *et al*. (2014) used SOLiD-based RNA-Seq approach to compare the expression of genes from three accessions of *Noccaea caerulescens* (Brassicaceae) with various abilities to tolerate and accumulate metals in order to identify genes linked to metal hyperaccumulation including nickel. In this study, the authors used the genome of the related model species *Arabidopsis thaliana* as a common reference to align RNA-Seq reads. Using an Illumina-based RNA-Seq approach, Meier *et al*. (2018) compared the expression of genes from various populations of *Senecio coronatus* (Asteraceae) hyperaccumulating (NiH) or not (NA) nickel. The authors generated a *S. coronatus* reference transcriptome by *de novo* assembly to quantify gene expression and identify differentially expressed genes in both type of populations. More recently, we used the same RNA-Seq technology to identify genes differentially expressed in pairs of NiH and closely related NA species from five distant plant families (García de la Torre *et al*., 2021). Then, to identify convergent mechanisms involved in nickel accumulation, we used orthologous relationship between genes from these distant families and a multiple testing correction to identify orthologous groups (OG) containing genes differentially expressed between NiH and NA species in at least 3 plant families.

These comparative approaches rely on the possibility to have access to pairs of closely related species or populations with contrasting capacity to accumulate metals in order to use a common reference sequence. However, the identification of such pairs of species is not possible in all plant clades. In addition, the output of these analyses strongly depends on the specific pair of species that have been selected for the study. Finally, none of these studies take into account the phylogenetic tree and the genetic drift associated with the selected species.

Phylogenetic relationships between species are known to induce correlations between trait measurements that can affect the analyses when ignored (Felsenstein, 1985). Phylogenetic Comparative Methods (PCMs) precisely aim at taking these relationships into account, and have been extensively studied over the last few decades (Harmon, 2019). In the context of gene expression, Bedford & Hartl (2009) studied several stochastic models, including the Ornstein-Uhlenbeck (OU) process. This process can be seen as modeling the evolution of a quantitative trait under stabilizing selection towards an optimal value (Hansen, 1997). Building on this process, Rohlfs and Nielsen (2015) proposed a phylogenetic ANOVA framework that takes into account both phylogenetic and individual variations. Individual variations represent both intra-specific variations and measurement errors, and ignoring them can lead to severe bias in PCMs (Silvestro *et al*., 2015; Cooper *et al*., 2016). This framework has been used to detect OG with significant mean expression shifts across groups of species from various clades of animal kingdom (Rohlfs and Nielsen, 2015; Stern and Crandall, 2018; Chen *et al*., 2019; Catalán *et al*., 2019).

In this work, we have implemented a methodology to identify orthologous groups (OG) with expression shifts linked to the nickel hyperaccumulation trait in a wide diversity of plant species. This method uses RNA-Seq datasets to produce reference transcriptomes by *de novo* assembly and quantify gene expression in each species. Genes from the different species are then associated to OG and the expression of OG identified in all species is then analyzed by PCM. Using this methodology, we have identified OGs previously associated with nickel hyperaccumulation as well as new candidate genes involved in this trait. We believe that this methodology can be used more generally to identify genes associated to complex traits in a wide diversity of species.

## Materials and Methods

### Collection of RNA-Seq datasets

RNA-Seq datasets used in this study were collected from NCBI bioprojects PRJNA476917 (García de la Torre *et al*., 2021), PRJNA312157 (Meier *et al*., 2018) and PRJNA657163. The selected samples correspond to RNA extracted from leaves of nickel hyperaccumulator (NiH) and non-accumulator (NA) species or accessions from the families Asteraceae, Brassicaceae, Cunoniaceae, Phyllanthaceae, Rubiaceae and Salicaceae (representing five orders of Eudicots) and sequenced with the Illumina HiSeq2000 paired-end sequencing technology. Information on the different samples is summarized in **Table 1**.

**Table 1.** Description of RNA-Seq samples

| Species | Families | Location | NiH | Short name | Bioprojects | SRA samples |
| --- | --- | --- | --- | --- | --- | --- |
| *Senecio coronatus*<br>Agnes mine | Asteraceae | South Africa | Yes | ScorA | PRJNA312157 | SRX1901460<br>SRX1901471 |
| *Senecio coronatus*<br>Kaapsehoop | Asteraceae | South Africa | Yes | ScorC | PRJNA312157 | SRX1901463<br>SRX1901464<br>SRX1901465 |
| *Senecio coronatus*<br>Galaxy mine | Asteraceae | South Africa | No | ScorB | PRJNA312157 | SRX1901479<br>SRX1901480<br>SRX1901481 |
| *Senecio coronatus*<br>Pullen Farm | Asteraceae | South Africa | No | ScorD | PRJNA312157 | SRX1901469<br>SRX1901470<br>SRX1901472 |
| *Noccaea caerulescens*<br>Firmiensis | Brassicaceae | France | Yes | Ncfi | PRJNA474900 | SRX4174673<br>SRX4174674 |
| *Noccaea montana* | Brassicaceae | France | No | Nmon | PRJNA474900 | SRX4174677<br>SRX4174678 |
| *Microthlaspi perfoliatum* | Brassicaceae | France | No | Mper | PRJNA657163 | SRX8947159<br>SRX8947160<br>SRX8947161 |
| *Geissois pruinosa* | Cunoniaceae | New Caledonia | Yes | Gpru | PRJNA476928 | SRX4261243<br>SRX4261244<br>SRX4261245 |
| *Geissois racemosa* | Cunoniaceae | New Caledonia | No | Grac | PRJNA476928 | SRX4261246<br>SRX4261247<br>SRX4261248 |
| *Phyllanthus luciliae* | Phyllanthaceae | New Caledonia | Yes | Phlu | PRJNA645979 | SRX8723728<br>SRX8723729<br>SRX8723730 |
| *Phyllanthus conjugatus* | Phyllanthaceae | New Caledonia | No | Phco | PRJNA645979 | SRX8723731<br>SRX8723732<br>SRX8723733 |
| *Psychotria grandis* | Rubiaceae | Cuba | Yes | Pgra | PRJNA476927 | SRX4261234<br>SRX4261235 |
| *Psychotria costivenia* | Rubiaceae | Cuba | Yes | Pcos | PRJNA476927 | SRX4261236<br>SRX4261237 |
| *Psychotria revoluta* | Rubiaceae | Cuba | No | Prev | PRJNA476927 | SRX4261238<br>SRX4261239 |
| *Psychotria gabriellae* | Rubiaceae | New Caledonia | Yes | Pgab | PRJNA476924 | SRX4261225<br>SRX4261226<br>SRX4261227 |
| *Psychotria semperflorens* | Rubiaceae | New Caledonia | No | Psem | PRJNA476924 | SRX4261228<br>SRX4261229<br>SRX4261230 |
| *Homalium kanaliense* | Salicaceae | New Caledonia | Yes | Hkan | PRJNA476925 | SRX4261218<br>SRX4261219<br>SRX4261220 |
| *Homalium betulifolium* | Salicaceae | New Caledonia | No | Hbet | PRJNA476925 | SRX4261221<br>SRX4261222<br>SRX4261223 |

### Transcriptome assembly and expression quantification

Several *de novo* assembled transcriptomes used in this study were previously published and available from the bioproject PRJNA476917 (García de la Torre *et al*., 2021). The transcriptomes of *Senecio coronatus* and *Microthlaspi perfoliatum* were assembled *de novo* with QIAGEN CLC Genomics Workbench v9 using the same assembly parameters (similarity ≥0.95; length fraction ≥0.75) as used in García de la Torre *et al*. (2021). For these assemblies, we used the paired-end Illumina reads SRX1901479 (*S. coronatus*), SRX8947157 and SRX8947158 (*M. perfoliatum*). For each species, the reads of each sample were mapped to the corresponding *de novo* transcriptome using CLC Genomics Workbench v9 (similarity ≥0.875; length fraction ≥0.75).

### Construction of ortholog group seeds and annotation

The sequence of predicted proteomes encoded by 12 plant genomes were downloaded from the PLAZA 4.0 Dicots database (Van Bel *et al*., 2018): *Arabidopsis thaliana* and *Brassica rapa* (Brassicaceae), *Gossypium raimondii* and *Theobroma cacao* (Malvaceae), *Carica papaya* (Caricaceae), *Prunus persica* (Rosaceae), *Cucumis melo* (Cucurbitaceae), *Glycine max* (Fabaceae), *Ricinus communis* (Euphorbiaceae), *Populus trichocarpa* (Salicaceae), *Solanum lycopersicum* (Solanaceae) and *Coffea canephora* (Rubiaceae). We used the meta-approach MARIO to build ortholog group (OG) seeds using the 12 proteome sequences (Pereira *et al*., 2014).

We performed the annotation of these OGs using the HMMER package (Eddy, 1998). For each OG resulting from the MARIO output, we first performed a multiple alignment with MUSCLE (Edgar, 2004) and created a HMM profile using hmmbuild. A profile database of the OG seeds was created using hmmpress. To annotate the OGs, we searched for the closest homologs in the SwissProt database with hmmsearch using the HMM profiles as queries. We extracted the function, EC numbers, and GO terms from the hmmsearch hits with the lowest e-value (e-value ≤ 10e-45) and transfered these annotations to the corresponding OGs.

### Assignment of contigs to Orthologous Groups

For each assembled transcriptome, we searched in each contig for the longest ORF on the forward and reverse strands and translated this ORF to obtain the putative encoded protein. The assignment of contigs to OGs was performed with the HMMER package (Eddy, 1998). We performed hmmscan using the translation of the longest ORF of each contig as a query against the OG seeds profile database. We assigned contigs to the OG profile having the lowest e-value (e-value ≤ 1e-10, coverage ≥20%).

### OG expression matrix construction

We built an OG expression matrix associating each OG to its level of expression in each sample. The expression level of an OG was calculated as the sum of the read counts corresponding to the contigs assigned to this OG in each sample. We also summed the lengths of all contigs assigned to a single OG to obtain an OG length matrix.

### Data normalization and transformation

We computed the normalization factors *n*_*i*_ for each sample *i* using the TMM method, implemented in the function calcNormFactor of the edgeR package (Robinson and Oshlack, 2010). To take the length of the OGs (sums of the lengths of each contigs within each OG) and the size of libraries into account, we then computed log2 RPKM [reads per kilobase per million reads, (Mortazavi *et al*., 2008)] using the following formula:

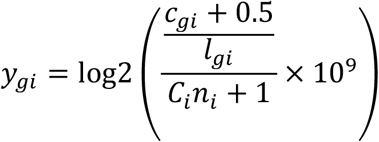

where 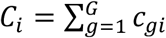 is the library size of sample *i, c*_”*i*_ is the read count for OG *g* in sample *i*, and *l*_”*i*_ is the length of the OG *g* in sample *i*. Note that the length can vary between samples for the same OG, as samples are taken from different species. To ensure that the ratio inside the log is strictly less than 1 and greater than zero, *c*_”*i*_ and *C*_*i*_*n*_*i*_ were offset away from zero by adding 0.5 and 1 respectively (Law *et al*., 2014).

To perform PCA on the OG expression matrix, we used the plot.PCA function of the R package

DESeq2. We used the rlog function to log-transform the data prior to the PCA.

### Phylogenetic tree

A custom dated phylogenetic tree was built from the plant tree backbone at the family level proposed by Magallón *et al*. (2015). The tree topology and time divergence used in our study are based on (Barrabé *et al*., 2014; Igea *et al*., 2015; Razafimandimbison *et al*., 2017) for Rubiaceae, (Pillon *et al*., 2014) for Cunoniaceae and (Huang *et al*., 2016) for Brassicaceae.

### Phylogenetic ANOVA

We used the R package phylolm (Ho and Ané, 2014) to perform phylogenetic ANOVA, using the “OU with fixed root” model, with measurement error. In a phylogenetic regression using an OU with fixed root model, the residuals are assumed to be correlated with a correlation between two species that depends on their shared evolutionary time (*ie* the time between the root of the tree and the most recent common ancestor of the two species). In addition to the phylogenetic residuals, we included in the model independent identically distributed residuals to capture additional non-phylogenetic variance, that we fitted to the data. The factor of interest was the nickel hyperaccumulation capacity of the species or accession. The design matrix also included an intercept, and a factor representing the country of origin of the sample. For each OG, a p-value was computed, corresponding to a t-test (with correlated observations) on the coefficient associated to the hyperaccumulator factor. A Benjamini-Hochberg multiple testing correction was applied to the vector of p-values. We used a threshold of 0.01, and selected OGs with a log2 fold change ≥ 1.5 or ≤ -1.5.

## Results

### A methodology for detection of orthologous genes with mean expression shifts in nickel hyperaccumulators from distant plant families

In this work we wanted to develop a methodology to identify genes whose expression is linked to nickel hyperaccumulation across a wide diversity of plant species (**Figure 1**). We took advantage of RNA-Seq datasets previously generated from nickel hyperaccumulator (NiH) and related non-accumulator (NA) species or populations to generate *de novo* transcriptome assemblies for each species and then use these transcriptomes as references to quantify gene expression for each sample corresponding to these species. This methodology also uses the concept of orthologous groups (OG) to annotate genes putatively playing conserved functions in distant plant families (Altenhoff *et al*., 2012). Finally, we quantified the expression of OG in NiH and NA species groups and analyzed the data with a Phylogenetic Comparative Method (PCM) to identify OG with an expression shift linked to the nickel hyperaccumulation trait.

**Figure 1:**
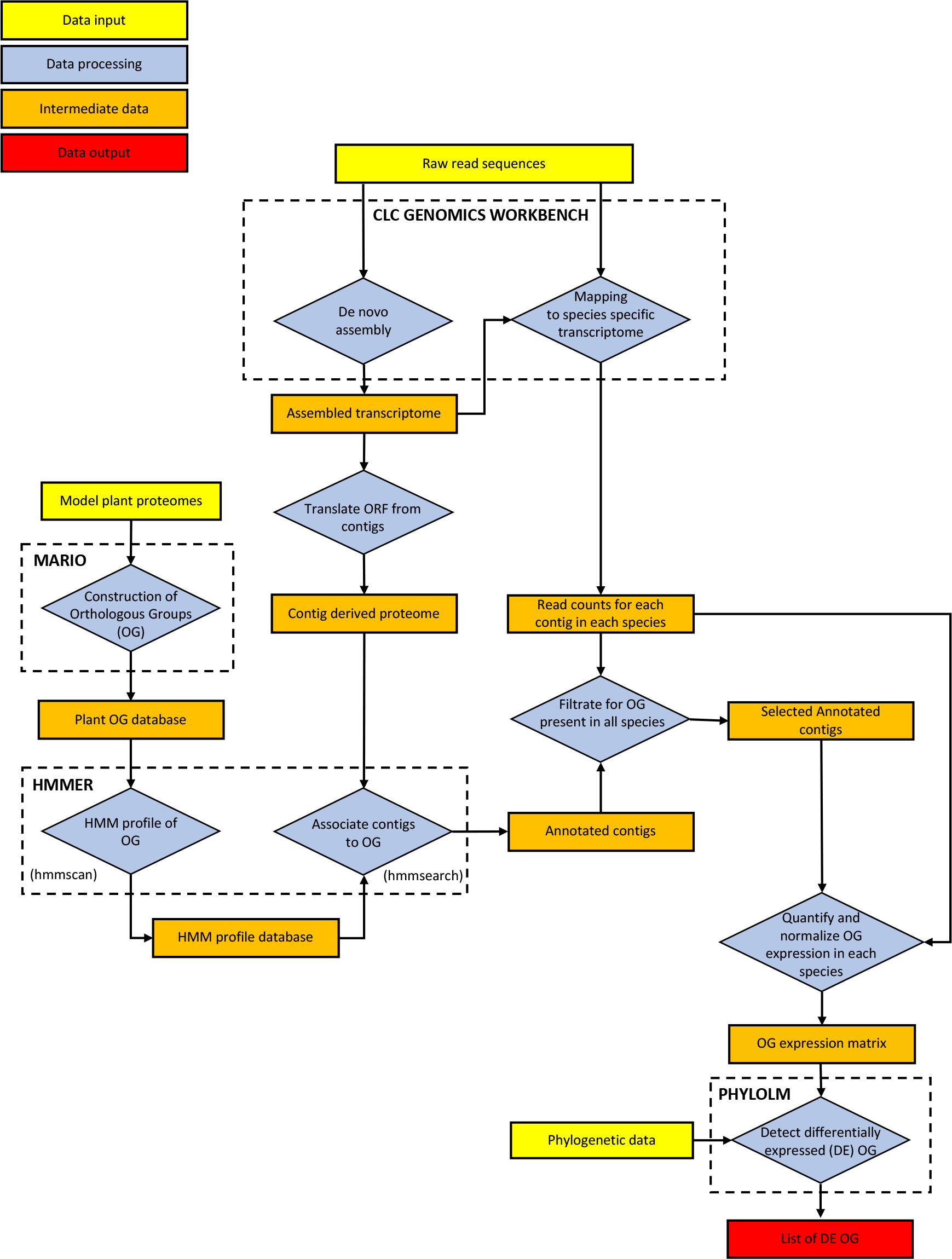
Workflow used to compare the expression of Orthologous groups in distant species (**a**) RNA-Seq datasets from different species were used to assemble transcriptomes *novo* and to quantify gene (contig) expression in these transcriptomes using QIAGEN LC Genomics Workbench. (**b)** Proteomes from sequenced plants species was used to generate a plant Orthologous group (OG) seed database using MARIO. (**c**) HMM profiles from the OG seed were produced by HMMER and used to annotate contigs from *de novo* transcriptomes. **(d)** The PHYLOLM package was used to detect differentially expressed OG using the normalized OG expression matrix.

### De novo transcriptome assembly and quantification of contig expression from nickel hyperaccumulator and non-accumulator species

We used available RNA-Seq from nickel hyperaccumulator (NiH) and non-accumulator (NA) species or accessions that were generated using Illumina paired-end technology (**Table 1**). For most of the NiH and NA species used in this study, the transcriptomes were previously assembled *de novo* using the CLC Genomic Workbench software (García de la Torre *et al*., 2021). We assembled the transcriptomes of *Senecio coronatus* and *Microthlaspi perfoliatum* using the same parameters. The number of contigs and the median size of the contigs for each species is given in **Table 2**. The number of contigs obtained for *M. perfoliatum* is significantly higher than for the other species probably due to the tetraploid nature of this species and because we assembled reads from both roots and shoots samples.

**Table 2.** *De novo* assemblies of contigs and assignment to OGs

| Species | Number of contigs | Contig median length | ORF median length | Number of classified contigs (%) |
| --- | --- | --- | --- | --- |
| Scor | 46726 | 475 | 330 | 27423 (59) |
| Ncfi | 41843 | 620 | 390 | 23608 (56) |
| Nmon | 64367* | 436 | 345 | 44460 (69) |
| Mper | 144815 | 420 | 312 | 89067 (62) |
| Gpru | 41188* | 486 | 288 | 20998 (51) |
| Grac | 37663* | 649 | 366 | 20410 (54) |
| Phlu | 42028 | 482 | 303 | 22136 (53) |
| Phco | 45524 | 437 | 288 | 22414 (49) |
| Pgra | 31362* | 728 | 456 | 19506 (62) |
| Pcos | 37309* | 542 | 321 | 19158 (51) |
| Prev | 45787* | 534 | 285 | 20254 (44) |
| Pgab | 46860* | 558 | 273 | 20493 (44) |
| Psem | 41854* | 600 | 285 | 19379 (46) |
| Hkan | 42325* | 541 | 300 | 22500 (53) |
| Hbet | 40774* | 537 | 306 | 21890 (54) |

\*contigs with low expression (TPM<1) were filtered out.

The RNA-Seq reads from each replicate corresponding to the leaf samples of all NiH and NA species or populations (**Table 1**) were mapped to the corresponding *de novo* transcriptome. This generated a read count table for each contig in all sample replicate from each species.

### Construction of orthologous group database and assignation of contigs

In the second step of our methodology, we wanted to establish orthologous relationships between genes expressed in the NiH and NA species in order to annotate genes potentially playing similar biological functions across these species (Altenhoff *et al*., 2012). However, because the translation of *de novo* assembled transcriptomes generates a large number of truncated peptides that could affect the analysis of the orthologous relationships, we decided to first build up an orthologous group (OG) library using proteomes predicted from sequenced plant genomes. We selected 12 sequenced plant genomes chosen along the dicotyledon phylogenetic tree and including species belonging to the same families as the nickel hyperaccumulators (see Methods). These proteomes, containing from 27000 to 56000 peptides, were used to create Orthologous Group (OG) seeds using the MARIO meta-approach (Pereira *et al*., 2014). MARIO combines the results of four methods: Best Reciprocal Hits, Inparanoid (O’Brien, 2004), OrthoFinder (Emms and Kelly, 2015) and Phylogeny (Lemoine *et al*., 2007) to establish orthologous relationships between proteins and compute a consensus OG annotation. Using this method, we obtained 17830 OGs containing from 2 to 1466 proteins. We could attribute a function to 11301 OG (63 %) using the Swissprot database and 4486 OGs (25%) were associated to at least one EC number. 7779 OGs (43%) are represented by at least one peptide in all model species. An HMM profile database was created from these OG seeds.

We then assigned each contig of the assembled transcriptomes from NiH and NA species to the OG seeds constructed with MARIO. For each contig, the longest Open Reading Frame (ORF) was identified and translated into a peptide. Each peptide was then associated to the closest OG using the HMM profile database (see Methods). Depending on the species, 44 to 69% of the contigs were assigned to an OG (**Table 2**). In total, among the 17830 OGs generated with the proteome of model plants, 15941 OGs are represented by at least one contig from at least one plant species of interest. It is important to note here that the RNA-Seq data used in this study only represent gene expressed in leaves. More importantly, for cross-species comparison, 5953 OGs are represented by at least one contig in each studied species.

### Identification of differentially expressed Ortholog Groups between nickel hyperaccumulator and non-accumulator species

We focused our differential expression analysis on the 5953 OGs represented in all plant species. We built an OG expression matrix (see Methods), each row representing one of the 5953 OGs and each column representing an RNA-Seq sample from 9 nickel hyperaccumulator (NiH) and 9 non-accumulator (NA) species or populations (2 to 3 biological replicates per species). We first performed principal component analysis (PCA) of the dataset after normalization and a log transformation of the expression data to explore the correlation between samples (**Figure 2**). The representation of the two principal components having the most important effect on total variation indicates that the RNA-Seq samples do not cluster with respect to the NiH trait but rather with respect to the plant family to which they belong. This result further illustrates that it is important to take into consideration the phylogenetic relationships between species in the transcriptomic comparison between the NiH and NA groups.

**Figure 2.**
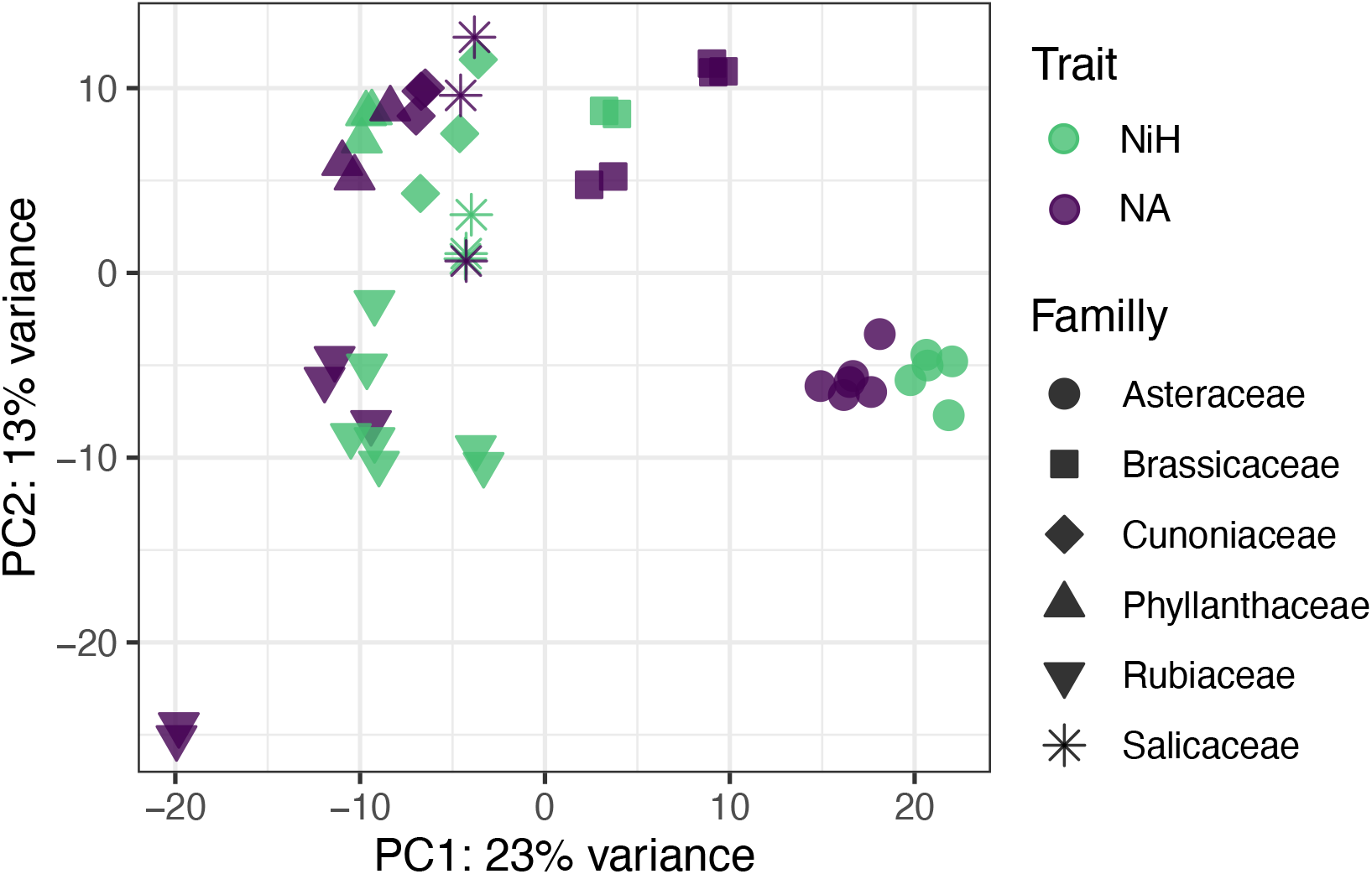
Principal Component Analysis performed on the OG expression table after rmalization and log transformation. The shapes of the symbols correspond to the family the species and the color to the nickel hyperaccumulation (NiH, green) or non-cumulating (NA, purple) trait.

To identify OGs differentially expressed between NiH and NA species from distant plant families, we used a phylogenetic mixed model (Lynch, 1991; Housworth et al., 2004) implemented in the R package phylolm (Ho and Ané, 2014).

Using this approach, we identified 31 OGs differentially expressed between NiH and NA species with a log2FC threshold ≥ 1.5 or ≤ -1.5. and an FDR ≤ 0.01 (**Table 3**). We used the length normalized unit RPKM to compare the expression of OG between samples. We also performed the analysis with the TPM unit (transcripts per million) (Wagner *et al*., 2012), but the results did not differ significantly (data not shown). 17 OGs are more expressed in the group of NiH species. The expression in NiH and NA species of OG 4147 and OG 7137, coding for a histidinol dehydrogenase and a membrane transporter of the SLC40A family respectively, is presented in **Figure 3**. These data illustrate the higher expression of these OG in several NiH species or populations compared to NA species. In contrast, 14 OG are less expressed in NiH species including OGs coding for a calcium-transporting ATPase (OG 10076) and a probable receptor-like serine/threonine-protein kinase related to Lr10 (OG 13722).

**Table 3.**
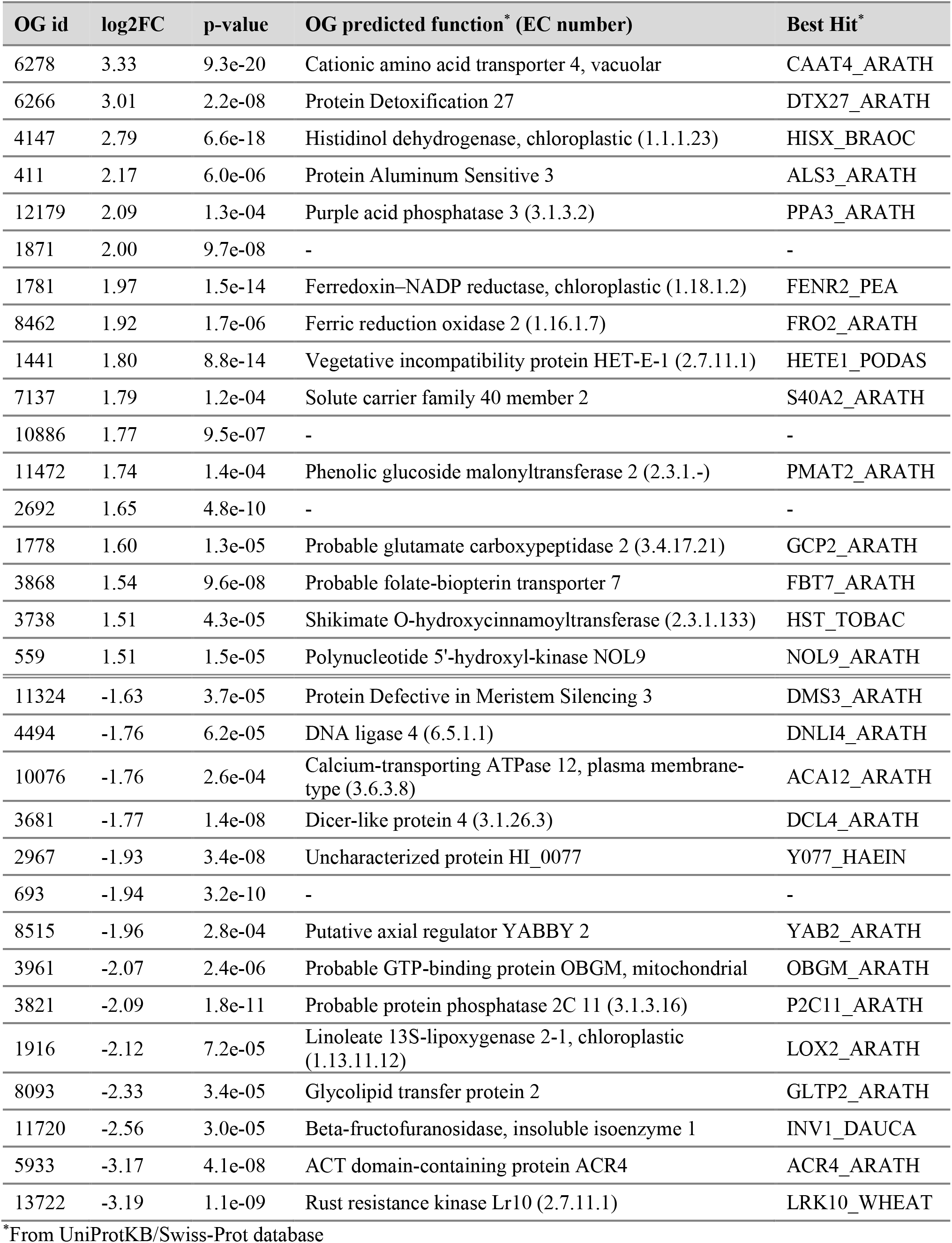
List of differentially expressed OG

| OG id | log2FC | p-value | OG predicted function* (EC number) | Best Hit* |
| --- | --- | --- | --- | --- |
| 6278 | 3.33 | 9.3e-20 | Cationic amino acid transporter 4, vacuolar | CAAT4_ARATH |
| 6266 | 3.01 | 2.2e-08 | Protein Detoxification 27 | DTX27_ARATH |
| 4147 | 2.79 | 6.6e-18 | Histidinol dehydrogenase, chloroplastic (1.1.1.23) | HISX_BRAOC |
| 411 | 2.17 | 6.0e-06 | Protein Aluminum Sensitive 3 | ALS3_ARATH |
| 12179 | 2.09 | 1.3e-04 | Purple acid phosphatase 3 (3.1.3.2) | PPA3_ARATH |
| 1871 | 2.00 | 9.7e-08 | - | - |
| 1781 | 1.97 | 1.5e-14 | Ferredoxin–NADP reductase, chloroplastic (1.18.1.2) | FENR2_PEA |
| 8462 | 1.92 | 1.7e-06 | Ferric reduction oxidase 2 (1.16.1.7) | FRO2_ARATH |
| 1441 | 1.80 | 8.8e-14 | Vegetative incompatibility protein HET-E-1 (2.7.11.1) | HETE1_PODAS |
| 7137 | 1.79 | 1.2e-04 | Solute carrier family 40 member 2 | S40A2_ARATH |
| 10886 | 1.77 | 9.5e-07 | - | - |
| 11472 | 1.74 | 1.4e-04 | Phenolic glucoside malonyltransferase 2 (2.3.1.-) | PMAT2_ARATH |
| 2692 | 1.65 | 4.8e-10 | - | - |
| 1778 | 1.60 | 1.3e-05 | Probable glutamate carboxypeptidase 2 (3.4.17.21) | GCP2_ARATH |
| 3868 | 1.54 | 9.6e-08 | Probable folate-biopterin transporter 7 | FBT7_ARATH |
| 3738 | 1.51 | 4.3e-05 | Shikimate O-hydroxycinnamoyltransferase (2.3.1.133) | HST_TOBAC |
| 559 | 1.51 | 1.5e-05 | Polynucleotide 5'-hydroxyl-kinase NOL9 | NOL9_ARATH |
| 11324 | -1.63 | 3.7e-05 | Protein Defective in Meristem Silencing 3 | DMS3_ARATH |
| 4494 | -1.76 | 6.2e-05 | DNA ligase 4 (6.5.1.1) | DNLI4_ARATH |
| 10076 | -1.76 | 2.6e-04 | Calcium-transporting ATPase 12, plasma membrane-type (3.6.3.8) | ACA12_ARATH |
| 3681 | -1.77 | 1.4e-08 | Dicer-like protein 4 (3.1.26.3) | DCL4_ARATH |
| 2967 | -1.93 | 3.4e-08 | Uncharacterized protein HI_0077 | Y077_HAEIN |
| 693 | -1.94 | 3.2e-10 | - | - |
| 8515 | -1.96 | 2.8e-04 | Putative axial regulator YABBY 2 | YAB2_ARATH |
| 3961 | -2.07 | 2.4e-06 | Probable GTP-binding protein OBGm, mitochondrial | OBGM_ARATH |
| 3821 | -2.09 | 1.8e-11 | Probable protein phosphatase 2C 11 (3.1.3.16) | P2C11_ARATH |
| 1916 | -2.12 | 7.2e-05 | Linoleate 13S-lipoxygenase 2-1, chloroplastic (1.13.11.12) | LOX2_ARATH |
| 8093 | -2.33 | 3.4e-05 | Glycolipid transfer protein 2 | GLTP2_ARATH |
| 11720 | -2.56 | 3.0e-05 | Beta-fructofuranosidase, insoluble isoenzyme 1 | INV1_DAUCA |
| 5933 | -3.17 | 4.1e-08 | ACT domain-containing protein ACR4 | ACR4_ARATH |
| 13722 | -3.19 | 1.1e-09 | Rust resistance kinase Lr10 (2.7.11.1) | LRK10_WHEAT |

\*From UniProtKB/Swiss-Prot database

**Figure 3.**
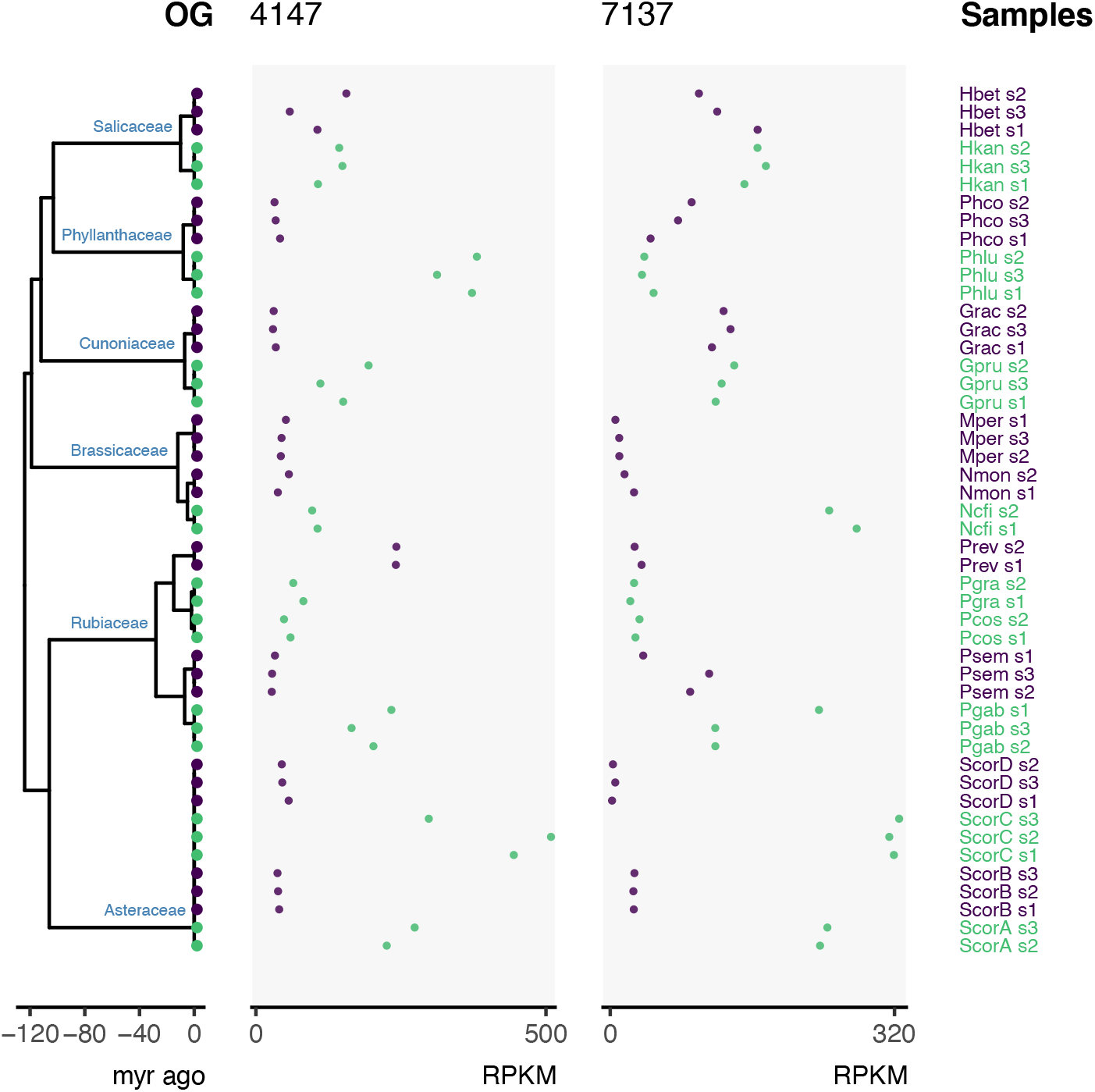
Representation of the expression of OG 4147 and OG 7137 along a plant ylogenetic tree. The expression of OG is expressed as Reads Per Kilobase of transcript, r Million mapped reads (RPKM). The phylogenetic tree is scaled in million years (myr), six families considered are labeled on the tree. The name of the samples is given on the ht. The color of the symbols correspond to NiH (green) or NA (purple) species or pulations.

## Discussion

The comparison of the expression of genes playing conserved function over a wide diversity of plant species is still a challenging task in comparative and evolution biology. The goal of this work was to implement a method using the annotation of orthologous groups (OG) and a Phylogenetic Comparative Method (PCM) to compare the transcriptomes of evolutionary distant species to identify genes whose expression level is linked with the nickel hyperaccumulation trait.

### Identification of orthologous genes with expression shifts linked to nickel hyperaccumulation

Genes linked to the nickel hyperaccumulation trait have been previously searched by comparing gene expression in pairs of closely related species or populations of the same species with contrasted capacity to accumulate nickel (Halimaa *et al*., 2014; Meier *et al*., 2018; García de la Torre *et al*., 2021; Enomoto *et al*., 2021). In this study, we have used available and comprehensive RNA-Seq datasets to identify OGs with expression shifts between nickel hyperaccumulator (NiH) and non-accumulator (NA) groups of species. Interestingly, we identified OG 7137 (**Table 3, Figure 3**) corresponding to the SLC40A membrane transporter family, also known as IREG or Ferroportin (FPN) transporters, as an OG more expressed in NiH compared to NA species. IREG/FPN transporters are able to transport divalent metal ions including nickel across membranes (Schaaf *et al*., 2006; Morrissey *et al*., 2009; Billesbølle *et al*., 2020). In *Arabidopsis thaliana*, two transporters belonging to OG 7137, AtIREG1 and AtIREG2, have been shown to localize on the plasma membrane and the vacuolar membrane respectively (Schaaf *et al*., 2006; Morrissey *et al*., 2009). High expression of vacuolar localized IREG transporters in transgenic *A. thaliana* increases nickel tolerance (Schaaf *et al*., 2006; Merlot *et al*., 2014). The high expression of IREG/FPN genes was previously shown to be linked to the nickel hyperaccumulation trait in species from several plant families (Meier *et al*., 2018; García de la Torre *et al*., 2021). The identification of OG 7137 thus validates the capacity of our methodology to identify OG linked to nickel hyperaccumulation. It also indicates that the high expression of IREG transporters in leaves is a robust characteristic of nickel hyperaccumulators that can be observed independently of the method used for cross species transcriptomic comparison. Our analysis also revealed a higher expression of OG 8462 and OG 6278, corresponding to the Ferric reductase FRO2 and the Cationic amino acid transporters CAT4 and CAT2 respectively, in the NiH species. These results confirm previous observations made in the Asteraceae species *S. coronatus* (Meier *et al*., 2018) and further suggest that the high expression of these genes corresponds to convergent mechanisms implicated in metal hyperaccumulation. Recently, the cationic amino acid transporter CAT4, primarily localizing on the vacuolar membrane, was associated with the histidine level trait in *A. thaliana* (Angelovici *et al*., 2017). Indeed, previous studies have highlighted the role of histidine as an important metal ligand involved in nickel hyperaccumulation in Brassicaceae species (Krämer *et al*., 1996; Kozhevnikova *et al*., 2014). Interestingly, our analysis also indicated that a high expression of Histidinol dehydrogenase (OG4147), the last step in histidine biosynthesis, is also associated with the nickel hyperaccumulation trait (**Table 3, Figure 3**). These results support a role for histidine synthesis and transport in nickel hyperaccumulation. The high expression of MATE and ABC transporters related to DTX27 (OG 6266) and ALS3 (OG 411), potentially involved in metal transport and tolerance (Larsen *et al*., 2004; Liu *et al*., 2009), was not previously associated with nickel hyperaccumulation. The validation of the contribution of these transporters to nickel hyperaccumulation will require further support.

On the contrary, our results suggest a lower expression of calcium-transporting ATPase (OG 10076) linked to the nickel hyperaccumulation trait. The rationale behind this result is not clear but it could be an indirect consequence of the edaphic conditions on which nickel hyperaccumulators are evolving. Indeed, NiH species are naturally growing on ultramafic soils that are rich in metals, including nickel, and with a strong calcium/magnesium imbalance affecting the development of most plant species (*aka* serpentine syndrome; Konečná *et al*., 2020). Therefore, the lower expression of calcium-transporting ATPase in NiH species might be linked to the adaptation of these species to the serpentine syndrome.

### Advantages and limitations of this methodology to compare distant species

One motivation of this work was to valorize the quantitative information contained in numerous RNA-Seq datasets already available from various plant species to identify gene functions associated with nickel hyperaccumulation. Most of these RNA-Seq datasets were produced with Illumina HiSeq paired-end technology. While the high number and quality of reads generated with this technology allow a good estimation of gene expression, their assembly frequently generates truncated ORF, thus affecting the annotation of OGs. To circumvent this limitation, we generated OG seeds using proteomes predicted from sequenced plant genomes. In a near future, the development of long-read sequencing technologies will undoubtably ease the assembly of full-length transcripts (Amarasinghe *et al*., 2020). In this work, we produced the OG database using the Mario method (Pereira *et al*., 2014). While this strategy allowed us to compare transcriptomes from distant species, the database is specific to this work and the results are thus difficult to compare with other studies. The use of more general OG databases for comparative and functional genomics such as Bgee for animals (Bastian *et al*., 2021) or PLAZA for plants (Van Bel *et al*., 2022), would thus favor comparisons between studies.

One main advantage of our method is that gene expression is quantified in each species using its own transcriptome. Therefore, this method does not absolutely require the identification of pairs of closely related species or populations with contrasted traits to quantify and compare gene expression using a common reference sequence. The identification of such pairs of closely related species with contrasted traits may represent a limiting condition in some genera. For example, the nickel hyperaccumulator *Blepharidium guatemalense* represents a monotypic genus (Navarrete Gutiérrez *et al*., 2021), and all species of the Cuban genus *Leucocroton* are able to hyperaccumulate nickel (Reeves *et al*., 1996). In addition, the output of pairwise comparisons to identify genes linked to a specific trait such as metal hyperaccumulation, strongly depends on the particular pair of species chosen for the comparison (Halimaa *et al*., 2014; Meier *et al*., 2018; García de la Torre *et al*., 2021). Our methodology based on the use of several species belonging to two contrasted phenotypic groups (*eg* NiH and NA) is less sensitive to the choice of species to identify conserved or convergent mechanisms linked in a complex trait.

The orthologous conjecture proposes that orthologous genes are functionally more similar than paralogous genes. While this conjecture is still disputed (Stamboulian *et al*., 2020), the annotation of groups of orthologous genes or orthologous groups is a widely used strategy for comparative analysis over a wide diversity of species (Van Bel *et al*., 2022). To compute the level of expression of OGs in each species, we decided to sum the counts of all contigs associated to the same OG as previously used (Lallemand *et al*., 2019). This choice is consistent with the hypothesis that contigs belonging to the same OG more likely encode for proteins playing the same function. However, OG may also contain inparalog genes, resulting from gene duplication in some species lineages, that might have acquired specific function. This is the case for OG 7137 containing for example *AtIREG1* and *AtIREG2* playing different roles in *A. thaliana* (see above). It is important to notice that this OG does not appear to be more expressed in NiH species from several families. This might be the consequence of the presence of inparalog genes in this OG with contrasted expression levels in species from some families, thus affecting the calculation of the OG expression level. Alternatively, the high expression of IREG in NiH species might not be a mechanism observed in NiH species from all plant families. A more detailed analysis of the expression of contigs composing this OG would be necessary to validate these hypotheses. However, the identification of OG 7137 suggested that the method using the sum of read counts does not prevent the identification of OG containing inparalog genes.

Altogether, our results suggest that this methodology based on the quantification of the expression of orthologous groups allows the identification of genes with expression shifts linked to nickel hyperaccumulation from distant plant species. Even though RNA-Seq sequencing technologies and comparative genomics resources are evolving rapidly, we believe that the methodology presented in this work could be used as a framework to identify genes linked to a specific complex trait in a wide diversity of plant species or other organisms.

## Data and code availability

The *Microthlaspi perfoliatum* Transcriptome Shotgun Assembly project has been deposited at DDBJ/ENA/GenBank under the accession GITW00000000. The version described in this paper is the first version, GITW01000000. The scripts and processed datasets are available at https://github.com/i2bc/plant-nickel-accumulation

## Contributions

M.G., O.L., S.M. conceived the study, V.S.GdlT., S.J., M.M., C.D., C.S., C.M, V.B. Y.P. collected the data, M.G., C.D., P.B., S.M performed the analyses and interpreted the results, M.G., O.L., S.M. wrote the manuscript. All authors have read and approved the final manuscript.

## Acknowledgement / Funding

The authors would like to thank Claire Toffano-Nioche (I2BC) for valuable discussions and support, and Bruno Fogliani (IAC/UNC) for his expertise on New Caledonian species. This work was supported by the X-TreM grant from the CNRS MITI to SM and the special funding MODELCOG from I2BC to MG. RNA-Seq sequencing for *Microthlaspi* and *Phyllanthus* species was financed by the ANR grant ANR-13-ADAP-0004 to SM, VB and BF.

